# Insufficient evidence for a severe bottleneck in humans during the Early to Middle Pleistocene transition

**DOI:** 10.1101/2024.10.21.619456

**Authors:** Trevor Cousins, Arun Durvasula

## Abstract

A recently proposed model suggests a strong bottleneck in the panmictic ancestral population of modern humans during the Early to Middle Pleistocene transition. Here, we show this model provides a substantially worse fit to the data than a panmictic model without the bottleneck.

## Main text

Recently, Hu et al. 2023 developed a new site frequency spectrum (SFS)-based approach, FitCoal, to infer effective population size changes in a panmictic population through time. The authors inferred a severe population bottleneck in the ancestors of modern humans approximately 1 million years ago (Mya) by applying FitCoal to present-day genome sequences from African populations. However, there are several issues with the analysis and conclusions. First, the authors did not evaluate how well their model fits the observed data relative to simpler models without a bottleneck. When performing demographic inference, it is important to recognize that the true history cannot be precisely determined. Instead, the principal goal of demographic history inference is to find the simplest models that can fit the observed data. New models that add complexity should only be preferred if they provide a better explanation to the observed data than simpler models. Without this crucial model comparison, it is unclear whether the model Hu et al. propose provides additional explanatory power. Second, FitCoal does not infer this bottleneck in out-of-Africa (OOA) populations, even though genetic events 1Mya should be shared by all present-day modern humans. Hu et al. argue this is due to OOA populations losing a substantial amount of genetic diversity after their ancestors left Africa ∼60kya, known as the OOA bottleneck, obscuring the ability to infer events in the deeper past. However, the amount of information about the ancient bottleneck available from the SFS of a population that experienced the OOA bottleneck remains unclear. Third, Hu et al. demonstrate with simulations that other methods, including PSMC (Li and Durbin 2011), Relate (Speidel et al. 2019), SMC++ (Terhorst, Kamm, and Song 2017), and Stairway Plot (Liu and Fu 2015), would have power to detect the bottleneck if it were truly there, but do not detect it in real data. This raises concerns about the robustness of the signal reported by Hu et al.

Here, we re-examine the findings of Hu et al. First, we perform formal model comparisons to test whether the bottleneck inferred by Hu et al. fits the data better than models without a bottleneck. Second, we test whether a bottleneck of the magnitude inferred by Hu et al. would leave a detectable signature in the SFS of OOA populations.

We processed data from the Yoruba (YRI) sequenced at high coverage by the 1000 Genomes project (Byrska-Bishop et al. 2022). We filtered out non-neutral regions of the genome (background selection statistic < 0.8 (Murphy et al. 2023)) and low-quality regions according to a mappability mask and generated an SFS. Using this SFS, we inferred an effective population size trajectory using FitCoal and saw a strong population bottleneck approximately 1 Mya, consistent with the results published by Hu et al. 2023 (**Figure 1A**). Next, we ran mushi (DeWitt et al. 2021) on the same SFS and found it did not replicate the finding of a strong bottleneck in the history of YRI (**Figure 1A**), echoing a similar recent failure to replicate the signal (Terhorst 2024).

**Figure 1.**
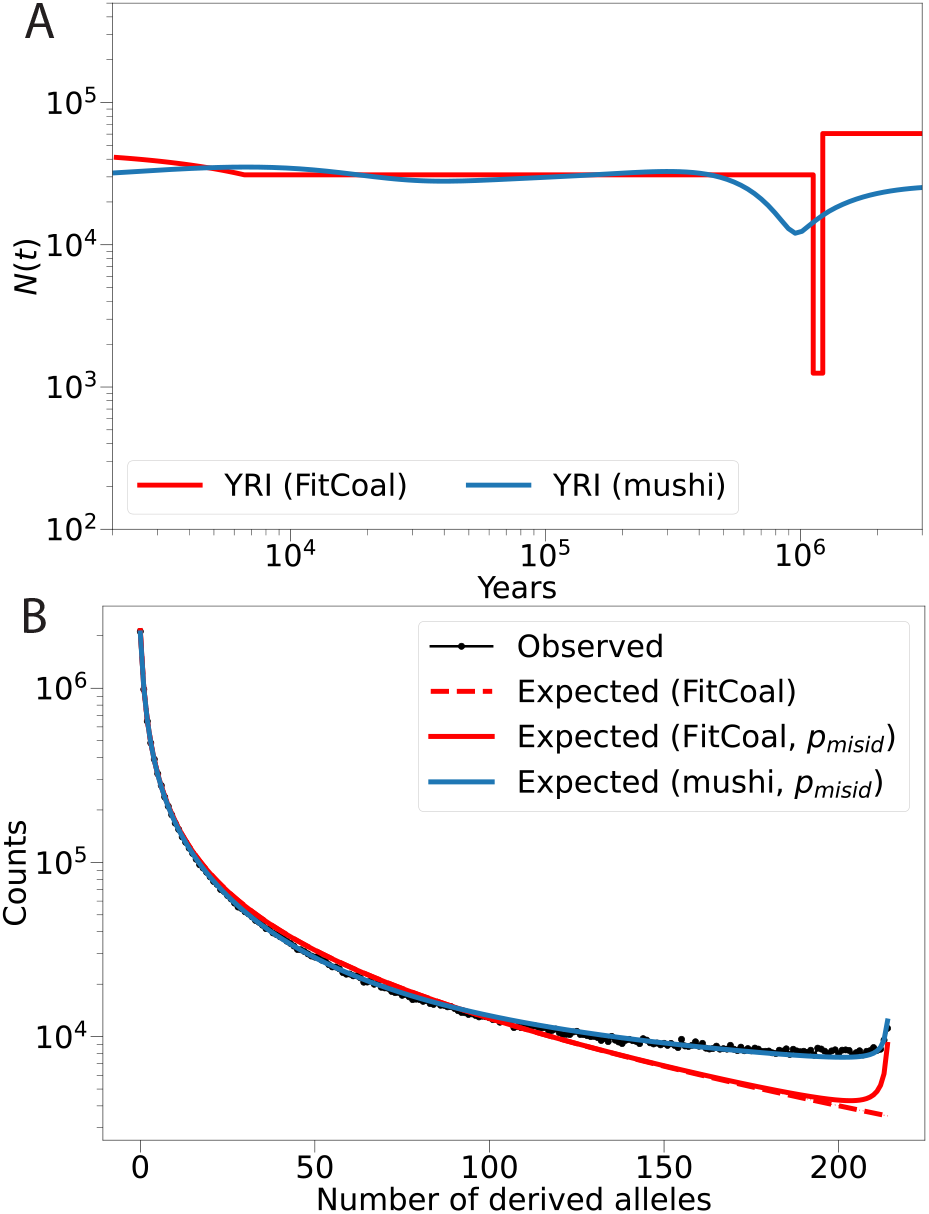
(A) Demographic histories inferred by FitCoal and mushi. (B) Site frequency spectra from the data (black) and models. *p*_*misid*_ uses the inferred ancestral allele misidentification parameter in mushi to accommodate the excess of high frequency derived alleles in the data 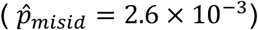. FitCoal does not fit an ancestral allele misidentification parameter. We used the value inferred by mushi to correct the expected FitCoal SFS.

The fact that FitCoal infers a bottleneck and mushi does not could reflect FitCoal’s sensitivity to ancient demographic history or alternatively statistical problems in the fitting procedure, as has been recently suggested (Deng, Nielsen, and Song 2024). Regardless, for a model to be favored over another model, it must fit the data better. We assessed this by computing the log-likelihood (*LL*) of the FitCoal and mushi models (**Figure 1B, Table 1**). We find the model fit by mushi provides an overwhelmingly better fit to the data (*LL*_*mushi*_ − *LL*_*FitCoal*_ = 69,903). This result suggests there is no reason to favor a model of panmictic human history with a severe bottleneck ∼1Mya compared to a panmictic model with no bottleneck.

**Table 1.** Model comparisons. We report the log-likelihood for the three models in **Figure 1** and difference in log-likelihoods (*ΔLL* with the best fitting model from mushi. *p*_*misid*_ refers to models that include an ancestral allele misidentification parameter.

| Model | Log-Likelihood ($LL$) | $\Delta LL$ with mushi model |
| --- | --- | --- |
| mushi ($p_{misid}$) | -1,503 | — |
| FitCoal ($p_{misid}$) | - 71,407 | 69,903 |
| FitCoal | - 92,937 | 91,433 |

Next, we investigated the discordance between the out-of-Africa (OOA) and African populations in the Pleistocene bottleneck. We computed the expected SFS of two demographic models: 1) both the Pleistocene bottleneck and the OOA bottleneck and 2) only the OOA bottleneck (**Figure 2A**). We find substantial differences between the SFS for each of these models (**Figure 2B**). This suggests a Pleistocene bottleneck of this magnitude would have left an identifiable signature in the genomic data of OOA populations, contrary to the claim of Hu et al. The fact that FitCoal did not infer a bottleneck in OOA populations raises further concerns about the robustness of the inference.

**Figure 2.**
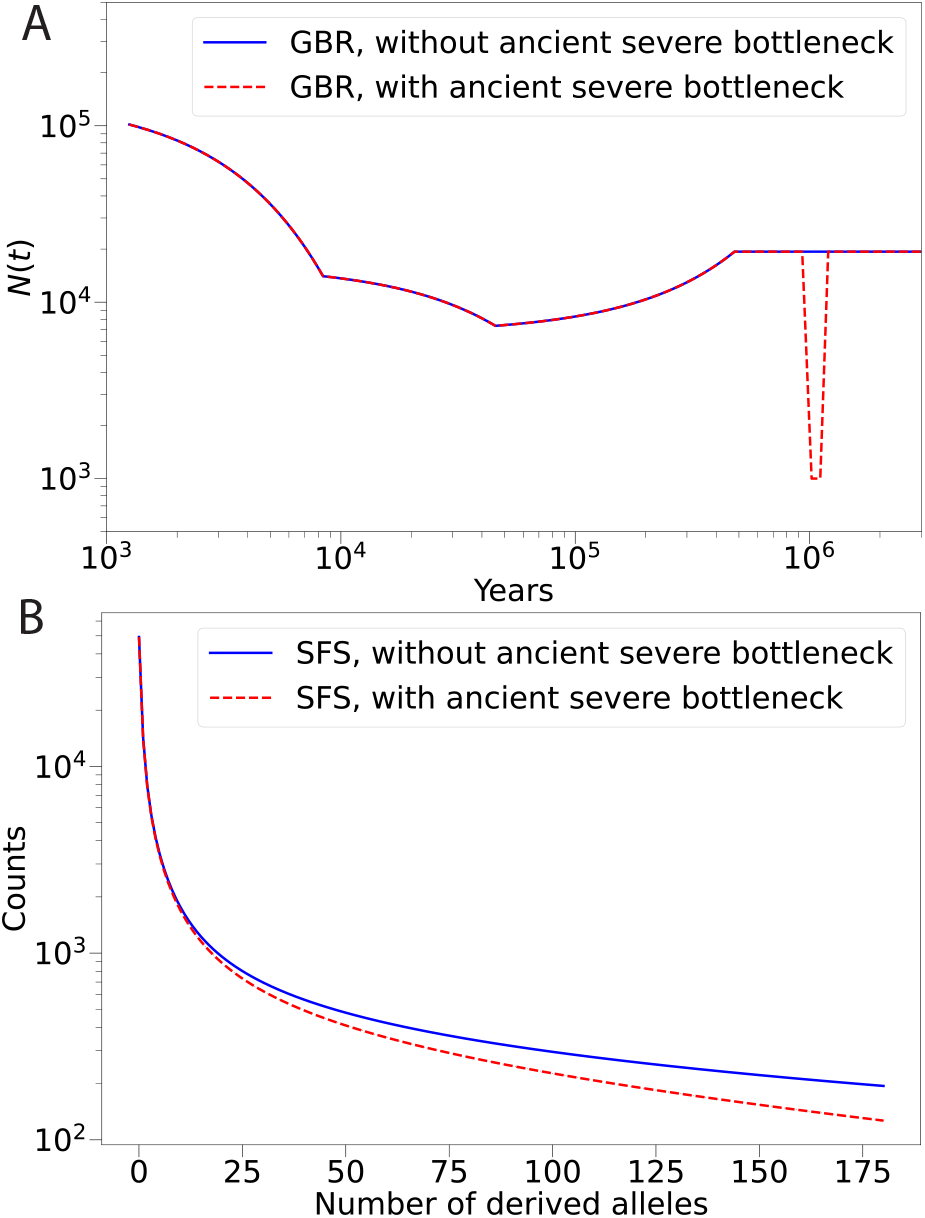
Effect of ancient severe bottleneck on site frequency spectra for out-of-Africa populations. (A) British (GBR) demographic history inferred by Hu et al. with FitCoal (blue) and the same demography with an added ancient severe bottleneck (red) (B) Expected SFS without (blue) and with (red) ancient severe bottleneck for out-of-Africa populations.

Finally, there is growing evidence that humans do not descend from a single, panmictic ancestral population. Instead, recent studies suggest that humans descend from two (or more) divergent populations that admixed together (Ragsdale and Gravel 2019; Lorente-Galdos et al. 2019; Durvasula and Sankararaman 2020; Wang et al. 2020; Ragsdale et al. 2023; Fan et al. 2023; Cousins, Scally, and Durbin 2024). These studies leverage different summaries of the genetic data, including linkage disequilibrium (Ragsdale and Gravel 2019; Ragsdale et al. 2023), the SFS (Lorente-Galdos et al. 2019; Durvasula and Sankararaman 2020; Fan et al. 2023), and heterozygosity across the genome (Wang et al. 2020; Cousins, Scally, and Durbin 2024), providing multiple lines of support for a structured evolutionary history. In particular, Cousins et al. 2024 proposed a model where modern humans descend from two ancestral populations that diverged 1.5Mya and admixed 300kya in a ratio of 80:20 percent, and that the majority ancestral population went through a bottleneck immediately after divergence. However, we do not believe this reflects the same event as inferred by Hu et al. using FitCoal. This is because methods used to infer population size history, including FitCoal, PSMC, Relate, SMC++, Stairway Plot, and mushi, estimate coalescence rates and assume the inverse of this rate reflects population size. While this assumption holds for panmictic populations, it does not in structured populations (Charlesworth 2009; Mazet et al. 2016). The coalescence rate in Cousins et al.’s model aligns with the estimates from PSMC, Relate, and SMC++, although the coalescence rate from Hu et al.’s model differs from all of these. Therefore, these bottleneck signals are not comparable.

In summary, we find there is insufficient evidence to support a model of human evolution where a panmictic population undergoes a severe bottleneck approximately 1Mya. We stress that newly proposed models should be shown to fit the data just as well or better than traditional models.

## Methods

### Processing SFS in real data

We downloaded variants aligned to GRCh38 from the Byrska-Bishop high coverage resequencing of 1KGP (Byrska-Bishop et al. 2022). We used the phased VCF from the publicly available data and excluded all positions with a B-value less than 0.8, based on the map of background selection from Murphy et al. 2023, which was lifted over from GRCh37 to GRCh38 using LiftOver (Hinrichs et al. 2006). Positions that could not be confidently lifted over were excluded from the analysis. We further removed positions that did not pass the strict mappability mask (ftp.1000genomes.ebi.ac.uk/vol1/ftp/data_collections/1000_genomes_project/working/20160622_genome_mask_GRCh38). To compute the SFS, we use the alternate allele counts at each position. We model ancestral allele misidentification using mushi (see below).

### Running FitCoal

We ran FitCoal (Hu et al. 2023) on the SFS data, and varied the number of high frequency SFS bins that are truncated, following Hu et al. We used truncation parameters of 40, 30, 26, 21, 20, 10, and 1 in all 26 1KGP populations. The truncation parameter did not significantly affect the results (as shown in Figure 1).

### Running mushi

We applied mushi (DeWitt et al. 2021) to the SFS data using the parameters trend_1 = 0, trend_2 = 1, and ridge = 1000. mushi includes a parameter for ancestral allele misidentification, 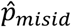, which was estimated as 2.6 × 10^−3^.

### Computing expected SFS

We used mushi to compute the expected SFS from the FitCoal model. We verified that mushi produces the correct SFS by running coalescent simulations of the same demography and generating an SFS using msprime (Kelleher, Etheridge, and McVean 2016).

### Evaluating model fits

The loglikelihood of the data was calculated by assuming each entry in the SFS is an independent Poisson variable (Sawyer and Hartl 1992; Gutenkunst et al. 2009). The loglikelihood was calculated as *LL* = ∏_*i*_ *o*_*i*_ In(*e*_*i*_) − *e*_*i*_ − *log*Γ(*o*_*i*_), where *i* indexes the number of derived alleles and Γ is the gamma function. To compute the likelihood of the FitCoal model with ancestral misidentification, we calculated the expected SFS as 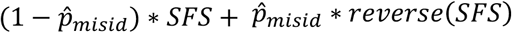

### Out-of-Africa bottleneck demography

We used mushi to compute the expected SFS for Out-of-Africa bottleneck models. The inferred demographic history was taken from Figure 3 of Hu et al. and downloaded from their Zenodo repository. To simulate a demography with an ancient bottleneck, we added a bottleneck of the same timing and magnitude as inferred in African populations by Hu et al.

## Acknowledgements

We thank David E. Reich, Nick Patterson, Regev Schweiger, and Aylwyn Scally for helpful discussions.

## Data availability statement

Variant call format files are available from https://www.internationalgenome.org/ and code to reproduce the analyses are available at https://github.com/trevorcousins/insufficient_evidence_panmictic_bottleneck/ along with processed data.

